# Controlling the frequency dynamics of homing gene drives for intermediate outcomes

**DOI:** 10.1101/2024.05.03.592448

**Authors:** Benjamin J. Camm, Alexandre Fournier-Level

**Affiliations:** School of BioSciences, The University of Melbourne, Parkville, VIC 3010, Australia

**Keywords:** CRISPR, modeling, simulation, conversion efficiency, selection, inbreeding, dominance, resistance, fitness cost

## Abstract

Gene drives have enormous potential for solving biological issues by forcing the spread of desired alleles through populations. However, to safeguard from the potentially irreversible consequences on natural populations, gene drives with intermediate outcomes that neither fixate nor get removed from the population are of outstanding interest.

To elucidate the conditions leading to intermediate gene drive frequency, a stochastic, individual allele-focused gene drive model accessible was developed to simulate the diffusion of a homing gene drive in a population. The frequencies of multiple alleles at a locus targeted by a gene drive were tracked under various scenarios. These explored the effect of gene drive conversion efficiency, strength and frequency of resistance alleles, presence and strength of a fitness cost for the gene drive, its dominance and the level of inbreeding.

Four outcomes were consistently observed: Fixation, Loss, Temporary and Equilibrium. The latter two are defined by the frequency of the gene drive peaking then crashing or plateauing, respectively. No single variable determined the outcome of a drive, instead requiring a combination of variables. The difference between the conversion efficiency and resistance level differentiated the Temporary and Equilibrium outcomes. The frequency dynamics of the gene drive within outcomes varied extensively, with different variables driving this dynamics between outcomes.

These simulation results highlight the possibility of fine-tuning gene drive outcomes and compensating through biotechnological design constraint imposed by population features. To that end, we provide a web application implementing our model which will guide the safer design of gene drives able to achieve a range of controllable outcome tailored to population management needs.

## Introduction

Gene drives are a promising, yet contentious, avenues of genetic biocontrol at our disposal to rectify pressing ecological issues. A gene drive can be used to alter, replace or suppress a population by spreading a desired allele through super-Mendelian inheritance (Gantz and Bier 2016; McFarlane *et al*. 2018; Rode *et al*. 2019). While gene drives have powerful features, they are equally hazardous. Gene drives can spread beyond their intended range (Greenbaum *et al*. 2021) or act in unintended ways (DiCarlo *et al*. 2015), potentially with detrimental consequences for the environment (Burt 2003). As such, modeling is a prudent approach to understand the key features determining gene drives’ behavior in a population before we can be confident that any synthetic gene drive will spread as intended (Combs *et al*. 2023). In particular, high confidence that a particular gene drive design will lead to the expected outcome is central (Prowse *et al*. 2017; James *et al*. 2020; Devos *et al*. 2022).

Previous models have explored how variation in pairs of variables can change either the outcome (Dhole *et al*. 2018; Rode *et al*. 2019) or the internal dynamics (Unckless *et al*. 2015; Dhole *et al*. 2020) of a gene drive. However, many of the features affecting a gene drive vary continuously, potentially interacting with each other to greatly complicate the outcome predictability (Nash *et al*. 2018; Frieß *et al*. 2023). Understanding how the many variables affecting a gene drive influence its fate thus requires their simultaneous testing over a continuous range of values. This would provide a more specific depiction of the effect each variable has on the frequency dynamics of a gene drive and finally its outcome.

Four distinct outcomes have been described for gene drives: the fixation of the gene drive allele, the loss of the gene drive allele (Gantz and Bier 2016), the transient establishment of the gene drive allele (Drury *et al*. 2017; Noble *et al*. 2019) or the gene drive allele frequency reaching an equilibrium within population (Unckless et al. 2015; Rode et al. 2019). The intermediate outcomes, either through transience or equilibrium, are often overlooked in gene drive models in favor of the more extreme fixation or loss outcomes when intending to suppress a population (Backus and Gross 2016; Paril and Phillips 2022). However, these outcomes could be of great relevance under specific scenario of population management. It may be safer, and arguably more ethical, to design a gene drive that is transient or reaches equilibrium in a population while the mechanics of the system is not yet fully controlled. A gene drive intended to suppress a local population may have unforeseen consequences that outweigh the potential benefits (Frieß *et al*. 2023). Designing gene drives that are locally self limiting over time and space are ideal candidates for initial gene drive releases (Marshall and Akbari 2018; Dhole *et al*. 2019, 2020).

The most commonly envisioned means to achieve self-limitation for a gene drive relies on geographic isolation, which requires the presence of reproductive barriers between target and non-target populations (Dhole *et al*. 2018; Greenbaum *et al*. 2021; Harris and Greenbaum 2023). Such spatial containment would rely on active monitoring and strict biosecurity measures that are difficult to enforce and costly (Giese *et al*. 2020). Alternatively, having a gene drive reach equilibrium in a population will avoid irreversible loss of alleles in the population, and potentially the whole species, and maintain genetic diversity at the gene drive locus. Given the inherent relevance of these intermediate gene drive outcomes, particularly in the short term, we need to understand which properties of a gene drive and the target population determine the outcome of a release, and how these properties affect the frequency dynamics over time.

Gene drive modeling has primarily investigated the end point of a gene drive (Unckless *et al*. 2015; Drury *et al*. 2017; Rode *et al*. 2019); mostly asking whether the gene drive would fixate or get lost. However, this also overlooks the many ways a gene drive can reach a given end point. If a gene drive fixates, does it fixate quickly or slowly? If a gene drive transiently reaches a maximum frequency, what is this maximum frequency? And then does the frequency remain stable or decrease? Understanding how the properties of a gene drive influence its dynamics will greatly increase the likelihood of meeting a desired outcome based on both population and biotechnological features. Furthermore, interactions between the properties of the gene drive and the target population could amplify or reduce their effects, and understanding how these properties interact as a complex system to influence the gene drive dynamics is crucial for safe gene drive design.

A range of gene drives properties have been tested as variables in models, from gene drive design (Dhole *et al*. 2018; Oberhofer *et al*. 2019; Noble *et al*. 2019), environmental effects (North *et al*. 2019) to organism biology (Sánchez C. *et al*. 2020). The effect of controllable variables, alone or in conjunction, on the frequency dynamics and outcome of a gene drive could be tuned by design or by management to achieve a desired results, potentially compensating for imponderable features of the biological system targeted. Gene drive risk assessment should be exhaustive across ecological contexts (Kim *et al*. 2023) as there will be no risk taken on large-scale exploratory trials (Baltzegar *et al*. 2018; Frieß *et al*. 2023). As technology is almost in hand, the challenge is to measure the importance of every feature to deliver eco-evolutionary relevant predictions for a gene drive release (Combs *et al*. 2023).

Our study intended to describe the variable space that leads to intermediate gene drive outcomes to guide safer, self-limiting gene drive designs (Marshall and Akbari 2018). By running simulations across a large variable space, we have narrowed down the combination of variables that lead to each other, and determined how these variable influence the frequency dynamics of the gene drive within a given outcome. Through this, we aim to better inform gene drive design to maximize the likelihood that a gene drive will safely and surely meet its intend aim.

## Methods

### Modeling Framework

A stochastic, allele-level CRISPR-based homing gene drive model implemented in the Julia programming language was developed to test the effect of homing conversion efficiency, resistance level, resistance frequency, selection coefficient (acting against the fitness cost of the gene drive), exposure to selection, dominance of and inbreeding (Table 1) on the frequency of a gene drive allele tracked over time in a population. The model cycles through discrete, non-overlapping generations, running through modules that simulate the life cycle of a sexually reproducing organism with discrete and non-overlapping transition between a pre-zygotic haploid and a post-zygotic diploid stage (Fig. 1a). The most important model information is captured in the vector of gene drive and wildtype allele frequencies *P*, which is used to compute the matrix of diploid genotype frequencies as the outer product of *P* with itself *G* = *P*⊗*_outer_ P*. The homing conversion, selection, genetic drift and migration alter the genotype frequencies are modeled as detailed below, using sequential modules computing the matrix of their effect on each genotype and updating the genotype frequency. Mutation is modeled by randomly sampling an allele of origin weighted by its frequency and adding the mutant allele to the vector *P* starting at frequency 1/2*n* where *n* is the population size and inheriting the other properties from the ancestral allele. Mutation and migration are set to zero in the simulations presented here.

**Table 1:** Variables definitions and distribution used in the Monte Carlo simulations.

| Variable | Definition | Distribution |
| --- | --- | --- |
| Conversion efficiency | Rate at which a gene drive allele is able to convert a wild type allele in heterozygous genotypes in one generation | $U(0,1)$ |
| Resistance level | Conversion rate of gene drive for resistant alleles | $U(0,1) \times$<br>Conversion |
| Resistance frequency | Initial frequency of alleles resistant to gene drive conversion | $U(0,1) \times 0.4$ |
| Selection coefficient | Selection coefficient of the drive | $U(0,1)$ |
| Exposure rate | Proportion of population exposed to selection acting negatively on gene drive heterozygous and homozygous genotypes | $U(0,1)$ |
| Fitness cost | Product of the Selection coefficient and Exposure rate | $U(0,1)$ |
| Dominance | Level of dominance of the gene drive allele on the fitness of heterozygous genotypes. 0: recessive; 0.5=additive; 1=dominant | $U(0,1)$ |
| Inbreeding | Reduction in frequency of heterozygous genotypes compared to the Hardy-Weinberg expectation | $U(0,1)$ |

**Fig. 1:**
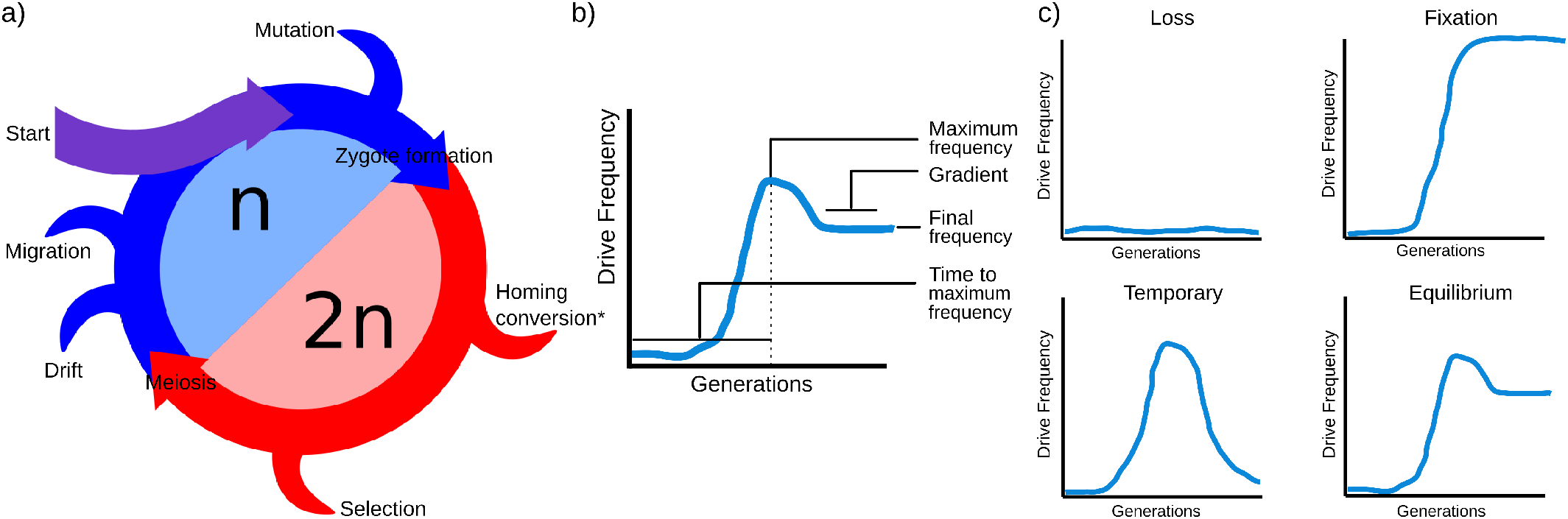
**a)** Model structure. The model is presented in the case of a postzygotic gene drive, prezygotic gene drives can be modeled by swapping the homing conversion and zygote formation modules. Mutation and migration are set to 0 in the simulations conducted in this study. **b)** Summary statistics used to define the categorical outcome of a gene drive release. **c)** Example s of frequency dynamics for each of the four outcomes.

### Gene Drive Homing Conversion

The conversion module simulates the CRISPR-mediated homing conversion of a wildtype allele into a gene drive allele in heterozygous individuals. Genotypes that are heterozygous for the gene drive allele have a probability to undergo conversion and become homozygous for the gene drive allele as:

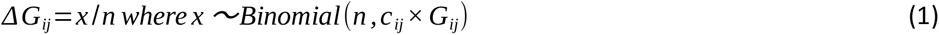

where *n* is the population size, *c_ij_* is the conversion efficiency of gene drive allele *i* on wildtype allele *j* and *Gij* is the frequency of the heterozygous genotype. The proportion of the heterozygous genotypes converted is subtracted from the frequency of wildtype alleles and added to that of the gene drive. For resistance alleles, the resistance level of the allele is substituted for the conversion efficiency. The resistance frequency value frequency of the resistance conversion efficiency allele in the population.

### Selection

The selection module simulates the effect of selection on diploid genotypes acting against the gene drive that assumes a fitness cost when exposed to selection. It factors three variables: the selection coefficient of the gene drive allele designed to model a fitness cost; the exposure rate which is the fraction of the population exposed to selection; and the dominance effect of the gene drive allele on the fitness cost of a genotype. Wildtype and resistant alleles have a fitness of 1. The fitness cost of a gene drive in a heterozygous genotype is affected by the dominance effect *d* of the gene drive allele, which ranges continuously from recessive (*d*=0), to additive (*d*=0.5) and to dominant (*d*=1). The survival of a genotype under selection, as determined by the exposure to selection, is proportional to the fitness of the genotype:

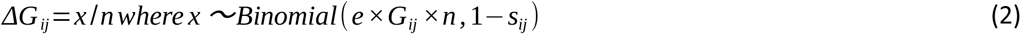

where *e* is the exposure rate of the population, *G_ij_*is the genotype frequency, *n* is the population size and *s_ij_* is the fitness cost of genotype *ij* equal to the product of the selection coefficient and exposure to selection. The change in genotype frequency (*ΔG_ij_*) is drawn and subtracted from the initial frequency. The *n* value is scaled by the percentage of the population exposed to the selection pressure. After selection, all genotypes are scaled to one.

### Drift

Drift is simulated by resampling each allele from a Binomial distribution and converting the outcome of a random draw divided by twice the population size into the post-drift frequency:

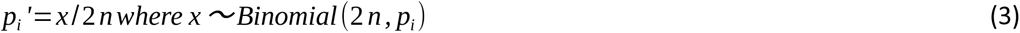

where *p_i_’* is the allele frequency of allele *i* after drift, *n* is the population size and *p_i_* is the initial allele frequency.

### Diploid zygote formation

Zygote formation leading to diploid genotypes assumes Hardy-Weinberg equilibrium except for allowing a level of inbreeding *F*, so that genotype frequencies for every combination of alleles *i* and *j* among *N* are given by:

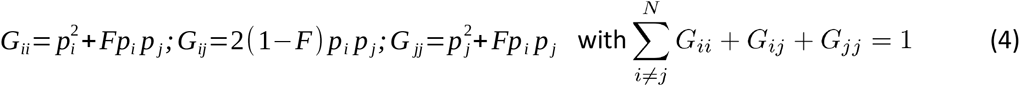

### Outcome classification

The outcomes were classified based on the maximum frequency, final frequency and change in frequency over time of the gene drive allele (Fig. 1b). The maximum frequency is the highest frequency the gene drive allele reaches over the course of a simulation. The final frequency is the frequency of the gene drive allele in the last generation. The change in frequency over time, referred to as gradient hereafter, is the change in gene drive allele frequency over the last 20 generations. A simulation was classified as Fixation if the final frequency is greater than 0.90; Loss if the maximum frequency is less than 0.10; Temporary if the maximum frequency is greater than 0.30 and a final frequency less than 0.10; Equilibrium if the final frequency is between 0.10 and 0.90 with the absolute value of th e gradient less than 0.0001 in the last twenty generations (Fig. 1c). Within these four gene drive outcomes, the allele frequency dynamics of the gene drive was analyzed using three metrics: the maximum frequency, the time to maximum frequency and the final frequency.

### Implementation

The modular structure of the model allows for the steps to be re-ordered to reflect different gene drive systems. This enables the modeling of both pre- and post-zygotic gene drive systems. In a post-zygotic gene drive system, conversion occurs before selection (Fig. 1a), whilst in a pre-zygotic system, conversion occurs after selection. Our stochastic model was validated against the results from published deterministic models (Suppl. Fig. S1-3; Unckless et al. 2015; Drury et al. 2017; Rode et al. 2019). Variation in our model was strictly due to random processes involving pseudo-random numerical sampling, and its magnitude primarily determined by the population size.

The model is available at adaptive-evolution.biosciences.unimelb.edu.au/SIEGE/ through a graphic user interface. Scripts to deploy and run the model locally can be sourced from https://github.com/bencamm001/interactive_gene_drive. A sensitivity analysis on the model variables was conducted using a multivariate Monte Carlo simulation framework. Variables in Table 1 were randomly sampled from a uniform distribution and the model was run 60,000 times to estimate the outcome frequencies (Table 2). In order to conduct further analysis with equal sample size across outcomes, another set of simulations was conducted, randomly sampling variables until 60,000 simulations of each outcome were observed. This was done to dissect the effect of all variable combinations across all outcomes with even power.

**Table 2:** Frequency of outcomes among 60,000 pre- and post-zygotic simulations.

| Outcome | Pre-zygotic Outcome Frequency | Post-zygotic Outcome Frequency |
| --- | --- | --- |
| Loss | 51.53% | 58.17% |
| Fixation | 33.65% | 32.02% |
| Temporary | 6.42% | 4.75% |
| Equilibrium | 1.34% | 0.42% |
| Total | 92.94% | 95.37% |

### Statistical analysis

Principal Component Analysis (PCA) was conducted using the Julia\MultivariateStats package (v0.8.0) on a down-sampled set of 2,500 simulations from each outcome to ease computation and visualization, using all seven variables listed in Table 1. Covariance ellipses for each outcome were drawn as elliptic contour line of a Gaussian density function with variance same variance as that observed for the data using the covellipse function from the StatsPlots package. This was done to qualify the importance of each variable in determining the outcome. Random Forest regression was used to measure the importance of all the variables in a random set of 40,000 simulations (10,000 from each outcome) from the sensitivity analysis using the R\randomForest package (v4.6.14). Variable importance was measured in terms of mean decrease in mean square error of each variable in determining a given outcome. Random Forest models were fitted to the data by drawing 500 trees using 3 variables for each tree. Random Forest was preferred over alternative modeling frameworks such as multivariate regression for its low sensitivity to overfitting and variable collinearity and ability to capture non-linear effects.

## Results

### Frequency of the different outcomes

A very high proportion of the 60,000 initial Monte Carlo simulations were classified as either Fixation, Loss, Transience or Equilibrium (95.37% for the post-zygotic and 92.94% for the pre-zygotic model). Under both models, Equilibrium was the rarest outcome, only found in 0.42% of post-zygotic simulations, and 1.34% of pre-zygotic simulations. The next rarest outcome was Transience, represented by 4.75% of post-zygotic simulations, and 6.42% of pre-zygotic simulations (Table 2). The uncategorized simulations corresponded to cases where the gene drive maximum frequency ranged between 0.1 and 0.3 (neither Loss nor Transience) or where the maximum frequency exceeded 0.1 but without plateauing (neither Fixation nor Equilibrium).

Principal component analysis (PCA) was used to assess how variables individually contributed to each outcome and how independent the different outcomes were over the variable space explored (Fig. 2a for pre-zygotic model and Fig. 2b for post-zygotic model). The analysis of post-zygotic gene drive simulations for each outcome showed that no single PC strongly distinguishing outcomes, which was qualitatively similar in pre-zygotic simulations. This suggested that no single variable was able to drive a specific outcome. Instead, the variable space covered by the different outcomes largely overlapped. Predictably, Loss and Fixation were the most distinct outcomes, with Temporary and Equilibrium representing intermediates between these. The variables with the strongest contribution to Fixation were the conversion efficiency and gene drive resistance, while the selection coefficient (modeling the fitness cost of the gene drive), exposure to selection and inbreeding level differentiated Loss from the other outcomes. The degree of dominance differentiated Temporary from the Equilibrium outcome. Interestingly, the resistance frequency had minimal contribution to outcome determination.

**Fig. 2:**
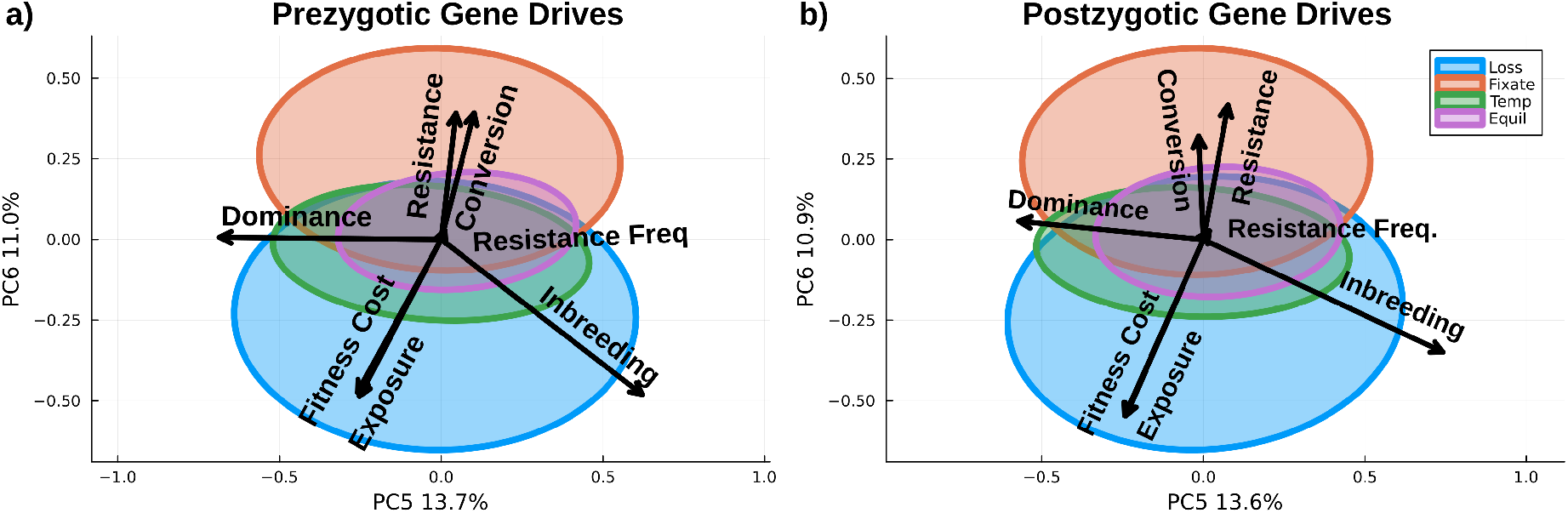
Principal Component Analysis of the post-zygotic Monte Carlo simulations based on the initial variables described in Table 1. PC5 and PC6 are used for a clear visualization of outcomes as all PCs explained a similar proportion of the variance. Covariance ellipses shown for clarity due to overlapping groups.

Variable importance measured through Random Forest (RF) models was consistent with the PCA. Dominance was the most important variable in determining the Equilibrium outcome with its non-inclusion in the model increasing the mean squared error by 365.16 (Table 3). Selection pressure (the product of selection coefficient and exposure rate) was important for all outcomes. Consistent with the PCA results, resistance frequency was not very important in driving outcomes overall, however, it was relatively important for the Temporary outcome. Both the PCA and RF regression results suggested that every variable tested could be used to an extent to influence the outcome, but it is likely that variables will need to be combined to efficiently determine the fate of a gene drive.

**Table 3:** Variable Importance on the outcome determination for a post-zygotic gene drive in Monte Carlo simulations measured in terms of mean squared error upon non-inclusion.

|  | Loss | Fixate | Temporary | Equilibrium | Full model |
| --- | --- | --- | --- | --- | --- |
| Conversion | 203.01 | 42.71 | 177.48 | 36.45 | 920.53 |
| Resistance | 69.02 | 307.75 | 339.3 | 311.6 | 1600.25 |
| Resistance Frequency | 7.99 | 2.38 | 40.69 | 19.35 | 299.41 |
| Selection Pressure | 277.22 | 385.46 | 320.89 | 337.08 | 2013.77 |
| Dominance | 105.02 | 42.58 | 104.58 | 365.16 | 1055.45 |
| Inbreeding | 297.73 | 151.84 | 245.1 | 312.35 | 1609.78 |

The distribution of variables within outcomes was used to assess the level of constraint on the value a given variable could take (Fig. 3), a narrow distribution indicating a limited space over which a variable can vary without modifying the outcome of the gene drive. Every variable except resistance frequency showed a constrained distribution for at least one outcome, with a maximum density skewed towards zero or one. The Loss and Temporary outcomes were constrained to resistance levels lower than 0.80, as opposed to Fixation and Equilibrium where the resistance level could take higher values (Fig. 3b). A similar pattern was observed for selection pressure (the product of selection coefficient and exposure rate; Fig. 3d) where Fixation or Temporary outcome were constrained to a selection pressure lesser than 0.5. Hence, targeting a Temporary outcome could only be achieved through the combination of both strong resistance level and weak selection pressure.

**Fig. 3:**
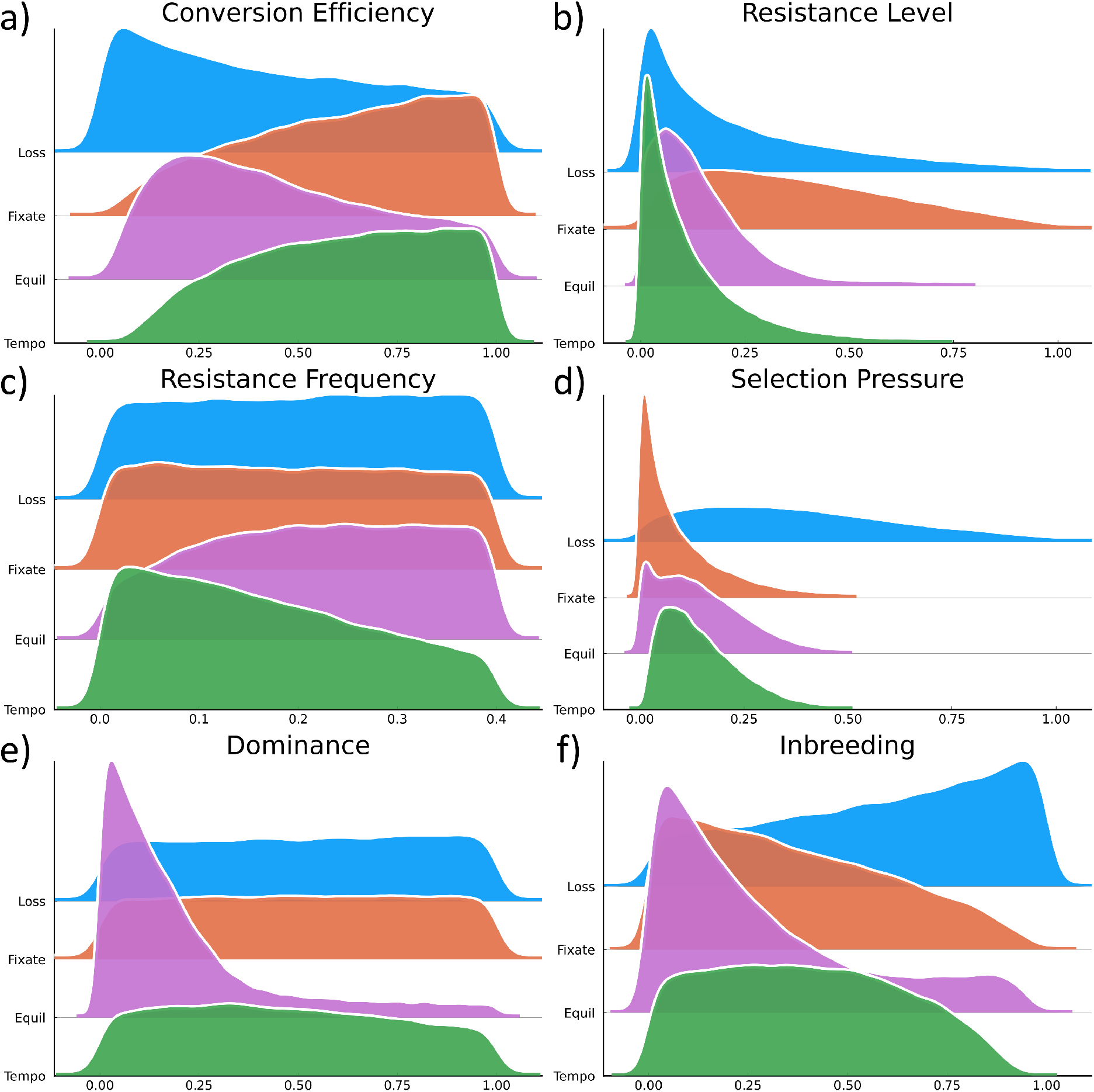
Distribution of the variables within each outcome. Resistance and selection pressure (the product of selection coefficient and exposure rate) show skewed distributions as they are the product of two uniform distributions.

The likelihood of an Equilibrium outcome dramatically increased for dominance value lesser than 0.5, requiring the fitness cost of the gene drive allele to be at best recessive or at best additive (Fig. 3b). Inbreeding less than 0.5 also promoted Equilibrium over other outcomes (Fig. 3f). A moderately low resistance level also promoted Equilibrium, but with less specificity (Fig. 3e). However, there was no bound variable space specifically promoting Equilibrium. This could be opposed to the Loss outcome that was frequently and specifically observed under high inbreeding or low conversion efficiency. Designing a gene drive for an Equilibrium outcome would thus require low levels of inbreeding and dominance, to which a non-null selection pressure should be added to avoid a Fixation or Temporary outcome. This level of constraint explains the rarity of the Equilibrium outcome across our simulations.

### Gene Drive frequency dynamics within outcomes

The Equilibrium outcome is inherently complex due to the broad variation in the gene drive maximum frequency and time to maximum frequency defining this outcome. Within Equilibrium outcomes, maximum frequency ranged from 0.1 to 0.99 and time to maximum frequency ranged from 16 to 500 generations (Fig. 4a). The simulations in which the time to maximum frequency extended towards 500 generations were characterized by high levels of inbreeding. Inbreeding slowed the increase in frequency of a gene drive to the point of equilibrium, so that the final frequency was also the maximum frequency. The final frequency of a gene drive at Equilibrium was most strongly determined by the resistance level and the selection pressure (Table 4). The final frequency was positively correlated to the resistance level (Fig. 4b) and negatively to the exposure rate (Fig. 4c). These two variables thus determined the equilibrium frequency of a gene drive. In turn, the time to equilibrium frequency depended on the conversion efficiency, with lower efficiency slowing the gene drive (Fig. 4d). Interestingly, the resistance frequency that had minimal importance in determining the outcome or the final frequency of the gene drive, yet had a strong effect on the time to maximum frequency (Table 4).

**Fig. 4:**
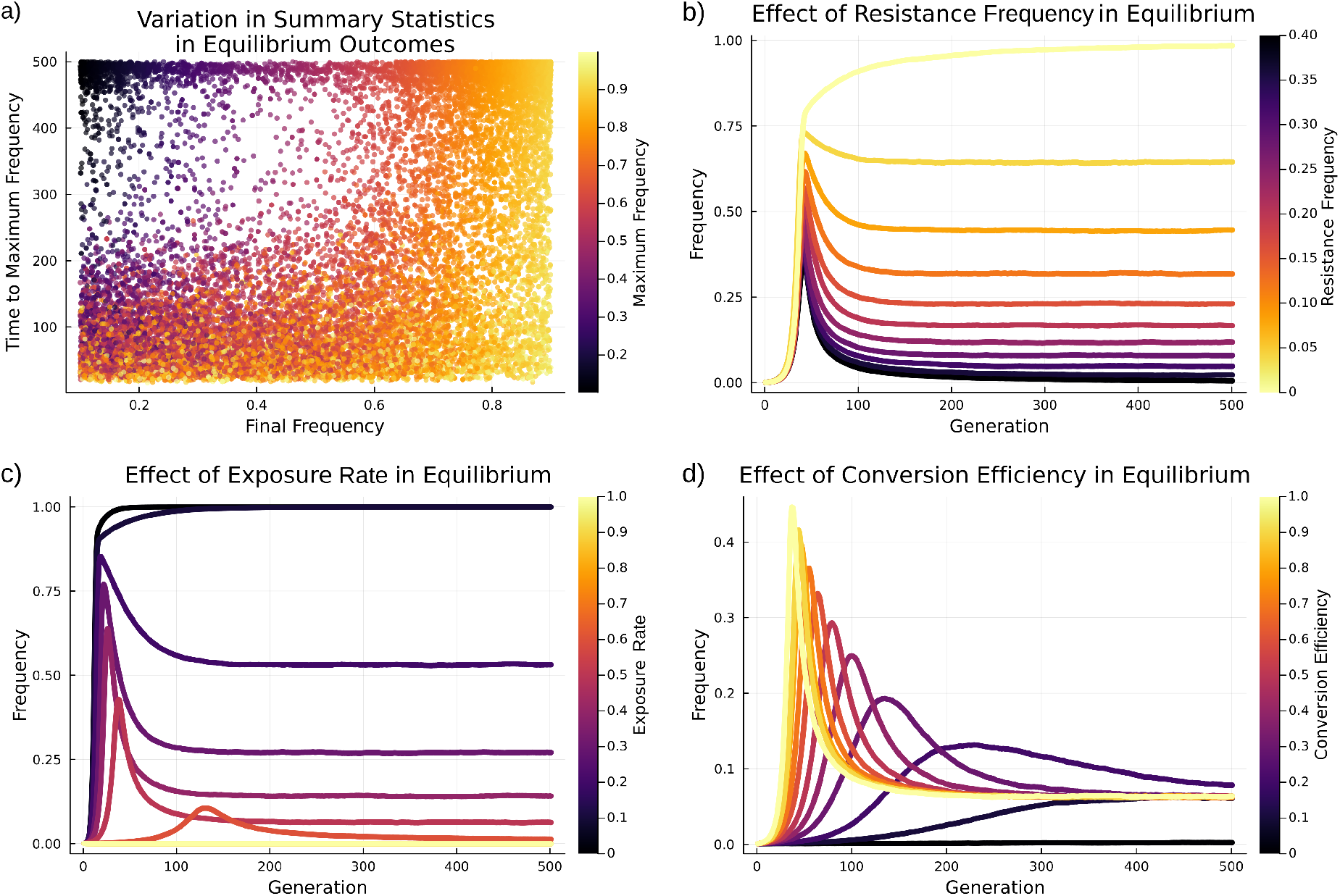
Effect of variables within Equilibrium outcomes. **a)** Variation in final frequency, maximum frequency and time to maximum frequency. **b)** Effect of wild type alleles resistance levels on gene drive frequency. **c)** Effect of exposure rate to selection against gene drive fitness cost. **d)** Effect of wild type conversion efficiency. Variables if not the focus of the analysis were set to: conversion efficiency = 0.95, selection coefficient = 0.8, exposure rate = 0.5, resistance level = 0.1, resistance frequency = 0.1, degree of dominance = 0.0 and inbreeding = 0.0.

**Table 4:** Effect of model variables on gene drive frequency within specific outcomes.

|  | Equilibrium outcomes |  |  |  |  |  | Temporary outcomes. |  |  |  |  |  |
| --- | --- | --- | --- | --- | --- | --- | --- | --- | --- | --- | --- | --- |
|  | Maximum frequency |  |  | Final frequency |  |  | Maximum frequency |  |  | Final frequency |  |  |
|  | Regression coefficient | Std. Error | Pr(>\|t\|) | Regression coefficient | Std. Error | Pr(>\|t\|) | Regression coefficient | Std. Error | Pr(>\|t\|) | Regression coefficient | Std. Error | Pr(>\|t\|) |
| Conversion efficiency | -276.72 | 2.3 | <1e-99 | -0.0203 | 0.0038 | <1e-7 | 0.216 | 0.00247 | <1e-99 | -195.92 | 0.79 | <1e-99 |
| Resistance level | 370.28 | 8.1 | <1e-99 | 0.6196 | 0.0134 | <1e-99 | -0.00772 | 0.00632 | 0.222 | 258.03 | 2.01 | <1e-99 |
| Resistance frequency | 302.67 | 5.04 | <1e-99 | -0.1307 | 0.0083 | <1e-55 | -1.22 | 0.00489 | <1e-99 | -29.97 | 1.56 | <1e-81 |
| Selection coefficient | -78.64 | 3.39 | <1e-99 | -0.4136 | 0.0056 | <1e-99 | -0.339 | 0.00352 | <1e-99 | 15.55 | 1.07 | <1e-47 |
| Exposure | -77.03 | 3.4 | <1e-99 | -0.4088 | 0.0056 | <1e-99 | -0.338 | 0.00336 | <1e-99 | 15.94 | 1.07 | <1e-49 |
| Dominance | 170.1 | 3.03 | <1e-99 | 0.1907 | 0.005 | <1e-99 | 0.055 | 0.00189 | <1e-99 | 18.94 | 0.6 | <1e-99 |
| Inbreeding | 240.09 | 3.12 | <1e-99 | 0.0215 | 0.0051 | <1e-04 | -0.229 | 0.00289 | <1e-99 | 147.17 | 0.92 | <1e-99 |

As with Equilibrium, Temporary outcomes could be defined for a broad range of maximum frequencies and times to maximum frequency. Within Temporary outcomes, maximum frequencies ranged from 0.30 to 0.99 and the time to maximum frequency ranged from 17 generations to 474 generations (Fig. 5a). The two variables showing the strongest effect on the maximum frequency within the set of simulations reaching a Temporary outcome were the resistance frequency and the selection pressure (Table 4). The initial frequency of resistant alleles was negatively correlated to the maximum frequency of the gene drive (Fig. 5b). This was due to the fact that the initial frequency of the resistance alleles, if remaining constant over time in the population, is the ceiling of the gene drive frequency. When the resistance frequency is zero, the gene drive fixates; when exposure of the fitness cost increases, the maximum frequency decreases (Fig. 5c). Contrary to what was found for Equilibrium outcomes, the time to maximum frequency in Temporary outcomes was most strongly regulated by the resistance level, followed by conversion efficiency (Table 4). Finally, increasing inbreeding and exposure rate both increased the time to maximum frequency, with higher levels of inbreeding also decreasing the overall maximum frequency of Temporary gene drives (Fig. 5d).

**Fig. 5:**
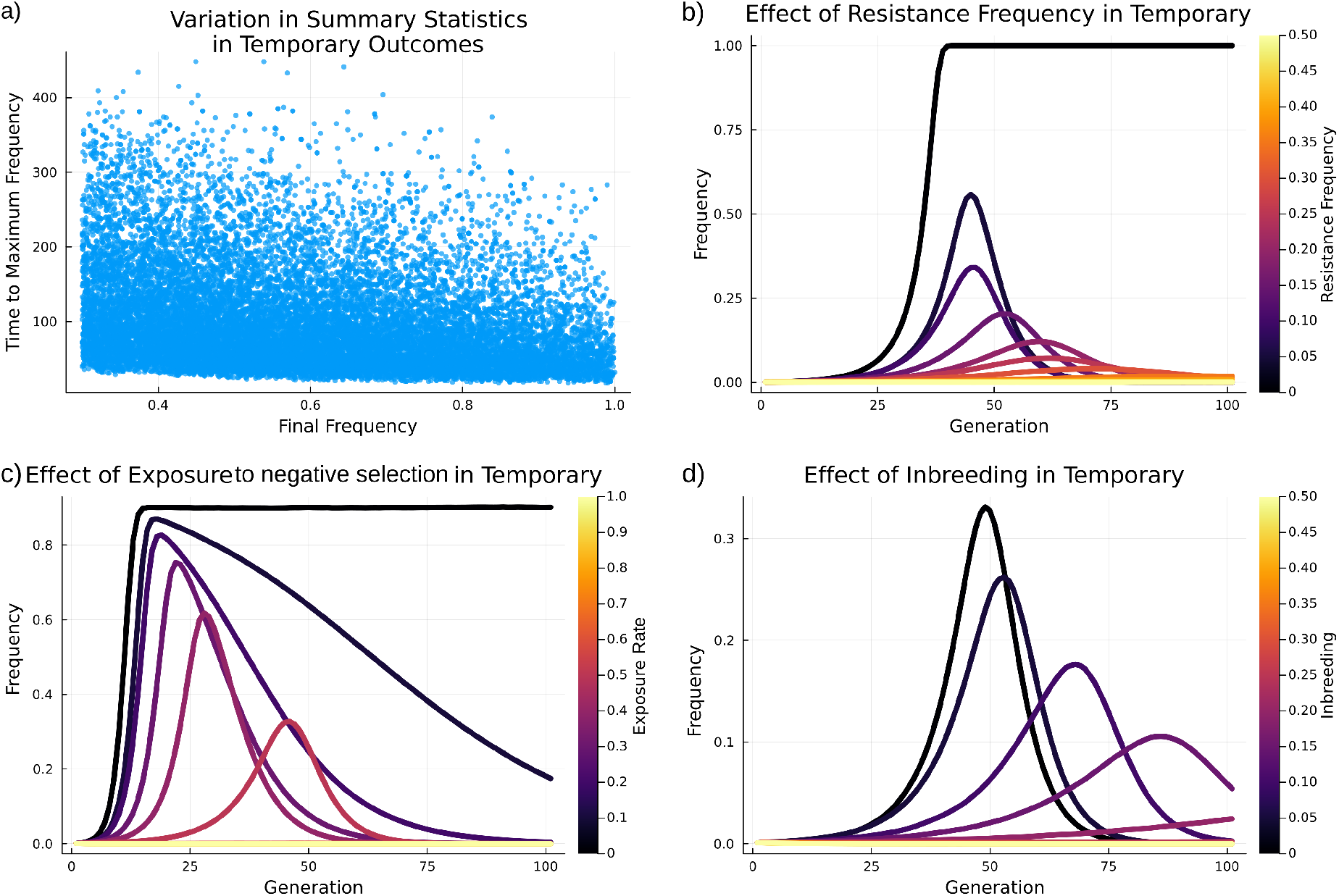
Effect of variables within Temporary outcomes. **a)** Variation in maximum frequency and time to maximum frequency. **b)** Effect of the resistance frequency. **c)** Effect of exposure rate to selection against gene drive fitness cost. **d)** The effect of inbreeding. Variables if not the focus of the analysis were set to: conversion efficiency = 0.95, selection coefficient = 0.8, exposure rate = 0.5, resistance level = 0.0, resistance frequency = 0.1, degree of dominance = 0.5 and inbreeding = 0.0.

### Switching from an Equilibrium to a Transient outcome

To assess the sensitivity of the gene drive frequency to the effect of a change in variables important determining outcomes, we investigated how a change in variable value during the simulation was able to change the outcome of a gene drive. For specific combinations of variable values, increasing the degree of dominance above 0.5 was able to shift the fate of a gene drive from Equilibrium, to Temporary to Fixation (Fig. 6a). The degree of dominance was thus pivotal to shift the fate of a gene drive between these three outcomes. It was also shown how changing the resistance conversion efficiency or exposure rate pushes a simulation from Fixation, to Equilibrium, to Temporary (Figure 4b-c). Changing the ratio between the conversion efficiency and resistance level was able to switch Equilibrium into Temporary outcomes (Fig. 6b). The ratio between conversion efficiency and resistance level was greater in Temporary than in Equilibrium outcomes. This suggests that if a Temporary outcome is intended, the conversion efficiency should be high and resistance level low, while for an Equilibrium outcome, the ratio between conversion efficiency and resistance level should be close to one.

**Fig. 6:**
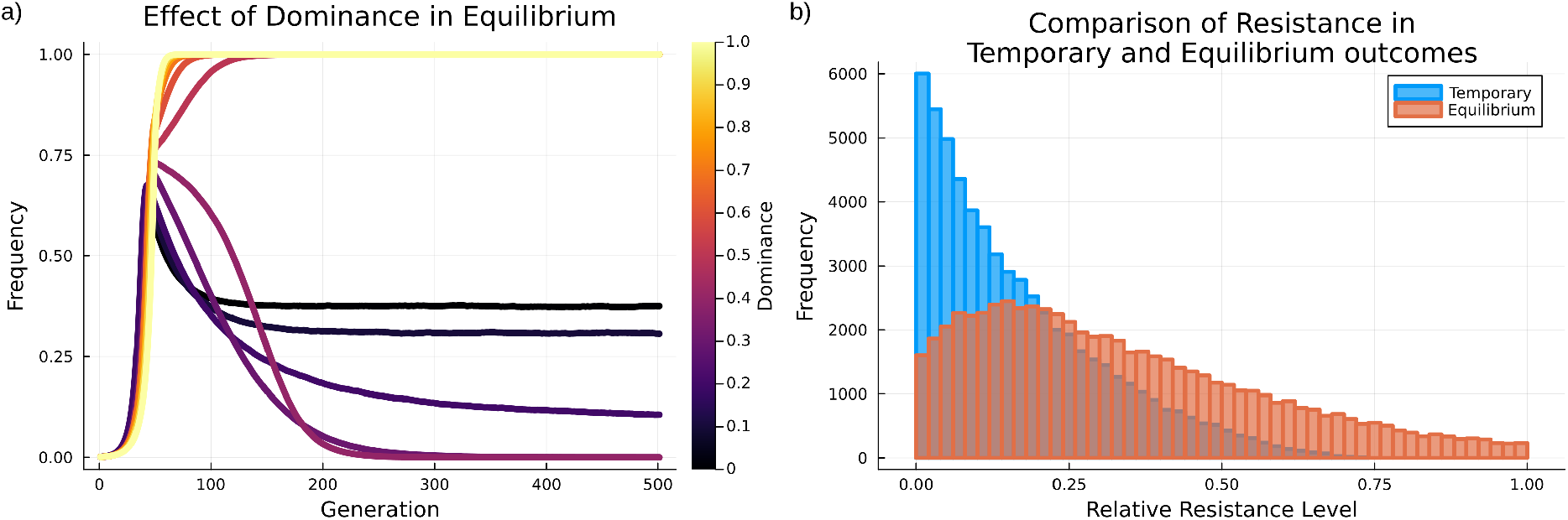
**a)** Effect of the level of dominance in promoting a switch between gene drive Fixation, Temporary and Equilibrium outcomes. Variables if not the focus of the analysis were set to: conversion efficiency = 0.95, selection coefficient = 0.8, exposure rate = 0.5, resistance level = 0.3, resistance frequency = 0.1 and inbreeding = 0.0.

## Discussion

Gene drives are complex systems influenced by multiple factors, some controllable by design, some set by the biology of the species or population targeted. Our work focused on the conditions able to promote intermediate Equilibrium and Temporary outcomes. While less frequent, they are particularly relevant for population management as they will not lead to irreversible loss of genetic diversity. These outcomes would be safer for early gene drive releases (Esvelt and Gemmell 2017; Noble *et al*. 2019), but required a combination of factors in our model. We identified specific combinations of values for the variables tested could lead to a given outcome, but also affect or alter the frequency dynamics of a gene drive to switch outcome.

### Balance between conversion efficiency and resistance for intermediate outcomes

For a homing gene drive to remain at intermediate frequency, the average conversion efficiency needs to be initially high before declining (Noble *et al*. 2019). The Temporary outcome can occur when the conversion efficiency differs between alleles in the population, or through the design of the gene drive. This can occur if the CRISPR elements lose over time their ability to convert other alleles into gene drives. This is likely to occur as no purifying selection is expected to maintain the integrity of the CRISPR elements. In contrary, a Temporary outcome can be achieved through engineered efficiency decay, as proposed in daisy chain (Noble *et al*. 2019) or split drive systems (Akbari *et al*. 2015; Kandul *et al*. 2020; Edgington *et al*. 2020). Alternatively, mutations in the gRNA target sequence of the gene drive may inhibit its function (Hsu *et al*. 2013; Unckless *et al*. 2017; Price *et al*. 2020; Grewelle *et al*. 2021). Genetic variation may segregate between populations, but also within population. Polymorphic alleles may be more or less complementary to the gRNA, leading to the type of quantitative resistance explored in our model (Willis and Burt 2021). Even with conversion efficiencies remaining constant, the frequency of the gene drive allele can increase, causing an increase in resistance allele frequency, and consequently reducing the gene drive allele frequency. The resistance frequency tends to impact the maximum frequency because it sets what fraction of the population is unlikely to be converted. The resistance level cannot be changed to influence the maximum frequency because as the resistance is lessened, it is less likely to result in a temporary outcome altogether. Gene drives often aim to reduce the carrier’s fitness, and conversion-resistant alleles expected to have a higher fitness, increase in frequency relative to the gene drive. As the resistant alleles increase in frequency through selection, the gene drive frequency will decrease until lost (Drury *et al*. 2017).

Equilibrium can rely on the balance between conversion efficiency and selection against the gene drive (Unckless et al. 2015; Rode et al. 2019), with the increase in gene drive frequency due to conversion matching the decrease in frequency due to its fitness cost. In absence of selection against the gene drive, a frequency equilibrium can be reached after all susceptible wild type alleles have been converted, only leaving fully resistant alleles behind. We show that such equilibrium is more likely under moderately efficient conversion. This is rather realistic as conversion efficiency is unlikely to be 100% in natural biological systems (Unckless *et al*. 2017; Price *et al*. 2020; Simoni *et al*. 2020). A gene drive designed to reach a stable, low frequency equilibrium would not fixate in the population and have less risk to escape to neighboring populations through gene flow. A gene drive that reduces the fecundity of *Plasmodium falciparum* could be designed to reach equilibrium in the population. This would not cause the extinction of the species but reduce the population size, possibly reducing their capacity for transmission, reducing the incidence of malaria (Tachibana *et al*. 2022).

### Constraints and feature Importance for designing gene drives

Consistent with previous studies, we determined that the variable space leading to intermediate outcomes is narrow (Eckhoff *et al*. 2017; Rode *et al*. 2019), with the strongest driver being the degree of dominance of the fitness cost. Random forest models identified rather accurately which aspect of the gene drive design to focus on to ensure a desired outcome. Ideally, an Equilibrium gene drive will have a recessive fitness cost and target an outbreeding population. However in practice, the level of control of each of the variable tested here varies greatly. The molecular properties of a gene drive can be controlled by design, some population features can be managed but others are intrinsic to the target population. For example, the level of inbreeding of the population is unlikely to be controlled, but the conversion efficiency, fitness cost and exposure rate could be adjusted by design.

Our analysis pointed at the critical importance of the balance between resistance level and conversion efficiency balance for controlling gene drive outcomes. Resistance to a gene drive can rise from multiple sources, whether standing genetic variation, *de novo* mutation or non-homologous end joining (NHEJ) after a failed gene drive homing (Price *et al*. 2020). Resistance to homing-based gene drives can be caused by mutations stopping the directed endonuclease protein from reaching its target. The likelihood of a mutation causing immunity can be limited by the use of Cas endonucleases edited to be protospacer adjacent motif (PAM) -promiscuous (Hu *et al*. 2018), or cut sites distal to the PAM site that are less likely to cause NHEJ (Paul and Montoya 2020). Gene drive resistance occurring through NHEJ after CRISPR cleavage can also be limited through gRNA multiplexing (Champer *et al*. 2020b).

We also showed that, as far as all wild type alleles are not fully immune to gene drive homing, it is possible to manage resistance. If the resistance allele has a low frequency, the gene drive is still able to increase in frequency up to a point of equilibrium with resistance alleles. At this point, if the gene drive frequency exceeds the equilibrium threshold, it will fixate; if it is below, it will become a Temporary drive and be lost (Unckless *et al*. 2015; Greenbaum *et al*. 2021). It is through this unstable equilibrium that a gene drive can overcome highly resistant alleles.

In a CRISPR-based gene drive, the similarity between the gRNA and its target site determines the cleavage efficiency, and therefore the conversion efficiency (Hsu *et al*. 2013). Hence, the resistance frequency can be controlled through gRNA target site choice in the same way the resistance level is controlled. Polymorphism in the gRNA target site can be leveraged to engineer gene drives to be Temporary or to reach Equilibrium. Future models will greatly gain in specificity and accuracy by incorporating information about the genetic diversity at gRNA target sites and its effect on the progression of a gene drive.

We found that gene drives are more likely to reach equilibrium in a population when acting pre-zygotically and when its fitness cost is recessive. While most of the modeling was done assuming a post-zygotic system, the effect of a pre-zygotic system was also explored. The pre-zygotic drive was more likely to create Temporary or Equilibrium gene drives, compared to the post-zygotic drive. This adds another layer of control over the outcome of a drive. The pre-zygotic gene drive system is more tolerable of high fitness costs, which may be useful for suppression gene drives, which are likely to confer a high fitness cost.

### Indirect control through exposure to natural selection

The frequency of an equilibrium can be fine-tuned when resistance and fitness cost are minimal and the control of resistance was previous discussed. The fitness cost can be manipulated, either by design or by varying the exposure to selection pressure through population management. Gene drives expressing a toxin are likely to be dominant and their cost will depend on the effect of the toxin (Champer *et al*. 2020a). For gene drives spreading susceptibility to a xenobiotic (Wedell *et al*. 2019; Bier 2022), the fitness cost will depend on the rate of exposure to selection. This exposure rate can be anthropogenically managed, through pesticide application for instance (Carrière *et al*. 2012). We have shown the pivotal importance of the dominance of the fitness cost for determining the outcome of a gene drive release. For gene drives targeting essential loci for which some level of genetic redundancy is expected, the fitness effect of the gene drive is likely to be more recessive. The way selection pressure is managed during the spread of a gene drive will thus be a critical aspect to manage to ensure the gene drive acts as intended by design.

A post-zygotic drive is unable to spread when selection against it is too strong, even under a recessive genetic architecture (Unckless *et al*. 2015). However, selection pressure is rarely homogeneous over time and space (Lenormand 2002; Carrière *et al*. 2012). Factoring an exposure rate in our model catered for the potential heterogeneity of selection that could be induced by, for example, a selection regime where a location where half the population is treated with a pesticide half is not. Modeling the ubiquitous application of a pesticide that kills 50% of the population is the same as applying a 100% lethal pesticide on half of a population. However, in a spatially explicit model, it may show changes in population structure due to the fitness differential. Most models have so far assumed that selection pressure is constant and ubiquitous when in most natural systems, selection varies over space and time. Relaxing selection through management practices (Karlsson Green *et al*. 2020) would allow the gene drive to spread in spite of a conditional high selection coefficient. This allows for more accurate modeling of a gene drive spreading susceptibility to a xenobiotic, such as herbicide resistance in weeds.

## Conclusion

Building confidence in the fact that gene drives will act as intended is vital. There has been too many dreadful examples of biocontrol measures becoming biosecurity issues (Blossey and Notzold 1995; Shine 2010; Schulz *et al*. 2019). Models provide guidelines and driving principles but can also be leveraged for the *in silico* stress-testing of new tools. These models will certainly be improved by incorporating more specific information about gene drives’ biology, such as how genetic variation within the gRNA target site may affect the conversion efficiency. Our work has explored how to design a gene drive that will achieve the intended outcome, but also for building confidence about our understanding of the frequency dynamics of a gene drive. This will aid in the development and design of future gene drives to maximize the likelihood that they will act as intended.

## Acknowledgments

BJC was supported by the CSIRO Synthetic Biology Future Science Platform.

## Data Availability Statement

Scripts for running the model are available at https://github.com/bencamm001/interactive_gene_drive, and can be accessed online at http://adaptive-evolution.biosciences.unimelb.edu.au/SIEGE/.

